# Engineered blue-shifted melanopsins for subcellular optogenetics

**DOI:** 10.1101/2023.10.07.561352

**Authors:** Dhanushan Wijayaratna, Filippo Sacchetta, Laura Pedraza Gonzalez, Francesca Fanelli, Tomohiro Sugihara, Mitsumasa Koyanagi, Senuri Piyawardana, Kiran Ghotra, Waruna Thotamune, Akihisa Terakita, Massimo Olivucci, Ajith Karunarathne

## Abstract

Melanopsin (MeOp) is a G protein-coupled Receptor (GPCR) family photopigment, expressed in intrinsically photosensitive retinal ganglion cells (ipRGCs) that display remarkable functional diversity. In addition to non-image-forming visual functions, MeOp also controls signaling underlying the retina development, circadian clock, mood, and behavior. MeOp is bistable, recycles retinal, and can function under low retinaldehyde availability. It also activates multiple G protein heterotrimers. Though MeOp could be a versatile optogenetic tool, its potential, especially its utility for subcellular signaling control, is hampered by the broader spectral sensitivity spanning the entire visible range. Here, we use a recently reported *in silico* technology called Automatic Rhodopsin Modeling (ARM) to identify blue-shifting mutations of MeOp and, ultimately, allow for imaging biosensors with red light without activating the opsin. Accordingly, ARM was used to construct validated quantum mechanics/molecular mechanics (QM/MM) models for mouse MeOp (mMeOp) to search and optimize a set of mutants featuring a blue-shifted light absorption. We demonstrate that four mutants of such can be successfully expressed and display the required resistance to activation by red light; however, they are activated by yellow, green, and blue light. Localized subcellular optical activation of these mutants in macrophage cells showed localized PIP3 generation and cell migration. Further characterization showed that MeOp blue-shifted mutants are also bistable. Altogether, our data demonstrate the computer-aided engineering feasibility of opsins with desired spectral properties for subcellular optogenetic applications.

## 1. Introduction

Optogenetics utilizes genetically encoded signaling actuators to control cell signaling and animal behavior using light^1-5^. Light-sensitive GPCRs (opsins) and their chimeric versions control G protein signaling and even animal behavior^6-10^. However, most opsins are monostable and thus become unstable after activation^11^. In the dark, a conserved *Lys* in the 7^th^ transmembrane (TM) region of opsins form a protonated Schiff base (PSB) with 11-*cis*-retinal (11CR) chromophore, and exposure to light induces 11CR-PSB isomerization to the all-*trans*-retinal (ATR)^12,13^; initiating the opsin photocycle. In the inactive dark state of monostable opsins, a negatively charged (i.e. deprotonated) *Glu* counterion residue on the 3^rd^ TM stabilizes the PSB via a salt bridge^14,15^. Upon light exposure, the PSB proton is transferred to the counterion, resulting in a large conformational change in the opsin, followed by the subsequent hydrolysis of the uncharged all-*trans* Schiff base^14-16^. This means that monostable opsins require a new 11CR molecule to reinstate photon sensitivity. Due to this continuous retinal demand, their optogenetic utility outside the retina is hindered, limiting *in vivo* applications of monostable opsins. In contrast, bistable opsins such as melanopsins (MeOps) utilize a *Glu* residue on the extracellular loop 2 (ECL2) above the retinal-binding pocket as the counterion for both 11CR and ATR. According to crystallographic studies, the *Glu* on ECL2 is too far from the Schiff base for a direct salt bridge formation, and therefore, a water-mediated hydrogen bond network makes Schiff base-counterion interactions possible^17^. Therefore, we hypothesized that bistable opsins are better candidates and provide potential templates for developing novel optogenetic tools since they can function under low 11CR concentrations^9^.

Most invertebrate opsins that evolved are bistable. Lamprey parapinopsin^18^ and Jumping Spider rhodopsin^17^ are examples. They evolved from the ancestral rhabdomeric photoreceptor cells. However, bistable vertebrate opsins such as MeOps exist. MeOp is expressed primarily in a subset of mammals’ retinal ganglion cells (RGCs), collects photic information from the retina, and sends it to the suprachiasmatic nucleus (SCN)^19^. SCN is the master circadian clock^20,24^. Furthermore, studies show that MeOp signaling affects mood and behavior, and, therefore, associated diseases, including seasonal affective disorder, depression, insomnia, and jet lag, have been linked to MeOp signaling^21^. MeOp is long been recognized as a Gq-coupled photopigment^22,23^. However, we and others showed that MeOp efficiently signals through Gq and Gi/o pathways, partly reasoning for its diverse functionality^24,25^. MeOp’s ability to activate additional G protein pathways has also been demonstrated^26^. Although these properties endowed MeOp with great potential as an optogenetic tool for live cell signaling control, its absorption spectrum with a λ_max_ of 440-480 nm made it sensitive to the entire visible range, preventing its use for controlling subcellular signaling^27,28^. Since subcellular optogenetic signaling control requires dedicated wavelengths for signaling activation and response imaging^29,30^, learning from blue opsin^6,29,30^, we hypothesized that the engineering of spectrally blue-shifted MeOp variants could, in principle, create MeOps that resist activation by red wavelengths.

Several studies examined shifting the absorption spectra of opsins, primarily to understand the evolution of color vision^31-34^. Previously, sequences of red and blue photopigments have been utilized to predict mutations that could spectrally shift the opsin absorption spectra^32,35-37^. However, we find that performing these studies using comparative structure-spectra analysis and best-guess approximations is experimentally inefficient and error-pronged. On the other hand, standard molecular modeling techniques based on molecular mechanics force fields cannot simulate light absorption and, therefore, could not assist such engineering efforts. This is because such force fields can only describe the dark state of the opsin but, due to the missing description of the retinal chromophore electronic structure, cannot describe the light-absorbing process. However, starting with the seminal work by Warshel^38^, the fields of computational photochemistry and photobiology have evolved to include hybrid quantum mechanical/molecular mechanical (QM/MM) calculations capable of systematically investigating and identifying spectral-shifting in wild-type (WT) and mutant opsins *in silico*^31,33,39^. These calculations account for the effects of chromophore-protein interactions, including electrostatic and steric effects on the stability of the electronic ground (S_0_) and excited state (S_1_) of the retinal chromophore^31,33,39,40^ and, therefore, on basic spectral properties such as the λ_max_; a quantity that can be easily calculated by converting the energy difference between the S_1_ and S_0_ states into the corresponding photon wavelength.

Despite their usefulness, the systematic construction and utilization of QM/MM models of WT and mutant opsins is far from trivial. Given the complexity of such models, this is a time-consuming and error-prone process. For these reasons, some authors were involved in developing a technology called Automatic Rhodopsin Modeling (ARM) for the automated construction of congruent opsin QM/MM models^41^. Here, we show the use of homology modeling and ARM, together with molecular engineering and live cell imaging approaches, to develop spectrally blue-shifted, bistable mMeOp mutants, and their applications in controlling subcellular signaling and cell behavior.

## 2. Results

### 2.1 Broad spectral sensitivity of melanopsin

MeOp has a λ_max_ between 440-480 nm^24,42,43^, and the uncertainty in the exact value is due to purification and spectral characterization difficulties. In order to determine the λ_max_ value, we expressed mMeOp with a carboxy-terminal 1D4 purification tag in HEK293S cells, purified the recombinant photopigment, and obtained the absorption spectrum as described previously^18,34,44^. Our experimental absorption maximum of MeOp-WT showed a λ_max_ of ∼470 nm (Fig. 1a). We then examined whether such a λ_max_ value makes mMeOp sensitive to blue as well as red wavelengths by examining mMeOp-induced activation of both Gq and Gi/o pathways (Fig. 1b)^25^. Gq pathway activates Phospholipase Cβ (PLCβ), inducing the hydrolysis of plasma membrane-bound phospholipid, phosphatidylinositol 4,5-bisphosphate (PIP2) into inositol triphosphate (IP3) and diacylglycerol (DAG). To measure the ability of blue, green, yellow, and red wavelengths to activate the mMeOp-Gq pathway, we employed PIP2 sensor (PH domain of PLCδ1) variants, each tagged with a different fluorescence protein, i.e., mTurquoise (mTq-434 nm), EGFP (484 nm), Venus (512 nm), or mCherry (587 nm), respectively (Fig. 1c). We expressed WT mMeOp and each PH sensor variant in HeLa cells. All experiments to examine the signaling of MeOp and its mutants henceforth were conducted in the presence of 10 μM 11CR, and cells were imaged only using the wavelength of consideration, unless otherwise specified. Imaging of the PIP2 sensor using 445, 488, 515, and 594 nm resulted in similar and robust PIP2 sensor translocation from the plasma membrane to the cytosol, indicating PIP2 hydrolysis due to the Gq pathway and subsequent PLCβ activation (Fig. 1c images and plot). We used 1.5 μW laser powers to image each fluorescence protein. The observed PIP2 hydrolysis attenuation that occurred during 300 seconds is due to a mechanism we have recently explained^45^, however, not the opsin deactivation.

**Figure 1.**
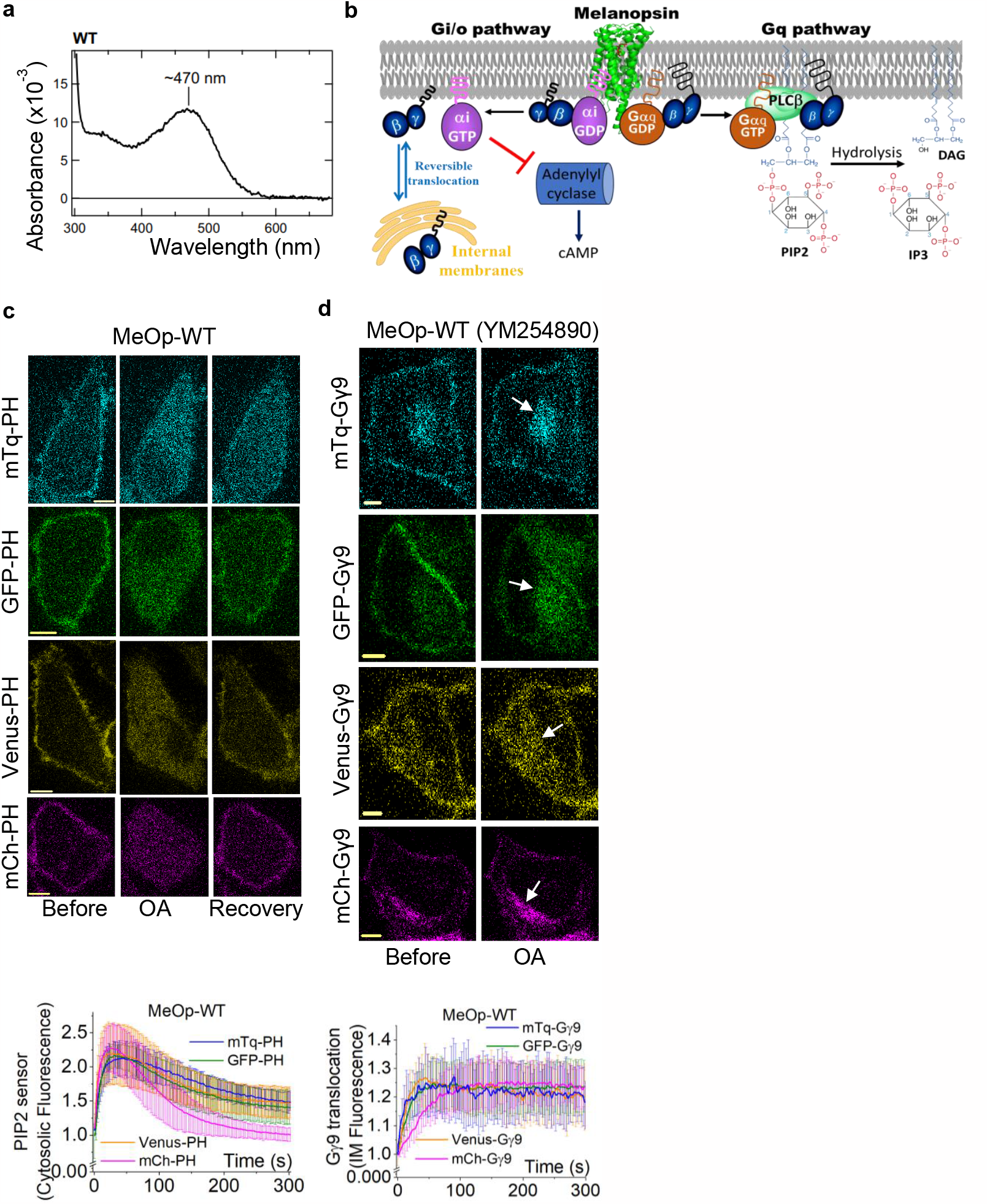
MeOp activates both Gq and Gi/o pathways upon sensing the entire visible spectrum. **(a)** The purified WT-MeOp photopigment exhibits an absorption maximum (λmax) of ∼470 nm. **(b)** Graphical representation of MeOp simultaneously activating both Gq and Gi/o pathways, which activate downstream signaling effectors. **(c)** Hela cells expressing WT-MeOp and PIP2 sensor exhibited robust PIP2 hydrolysis and subsequent recovery upon MeOp photoactivation by blue (445 nm), green (488 nm), yellow (515 nm), and red (594 nm) colors. Images show the PIP2 sensor tagged with fluorescent proteins of respective colors translocating from the PM to the cytosol upon MeOp activation. The corresponding plot shows the PIP2 sensor dynamics in the cytosol of the cells in response to different light colors. (n = 48) **(d)** HeLa cells exhibited robust Gγ9 translocation from the PM to internal membranes (IMs) upon MeOp activation by blue, green, yellow, and red light in the presence of 500 nM Gq inhibitor (YM254890). White arrows indicate the fluorescence intensity increase on IMs due to the Gγ9 translocation. The corresponding plot shows Gγ9 dynamics on the IMs. (n = 46). Average curves were plotted using ‘n’ cells, n = number of cells. Error bars represent SD (standard deviation). The scale bar = 5 μm.; MeOp: Melanopsin; WT: wild type; PLC: Phospholipase C; PIP2: phosphatidylinositol 4,5-bisphosphate; PM: Plasma membrane; IMs: Internal membranes; OA: optical activation

Next, we examined the Gi/o pathway activation by WT-MeOp in blue, green, yellow, and red wavelengths using the Gγ9 translocation assay we developed to measure G protein activations at the subcellular, single-cell, and multi-cell levels. This assay is based on the translocation of free Gβγ generated upon GPCR activation from the plasma membrane to internal membranes, significantly increasing internal membrane fluorescence^46,47^. Gγ9 has the most efficient translocation properties in the Gγ family^46,47^. To eliminate any influence from the Gq pathway to Gγ9 translocation, we performed the experiments in the presence of the Gq inhibitor, YM254890 (500 nM)^48^. Similar to the above Gq pathway assay, in the presence of 10 μM 11CR, we examined the translocation of mTq-Gγ9, GFP-Gγ9, Venus-Gγ9, or mCh-Gγ9 in HeLa cells also expressing mMeOp (Fig. 1d images and plot), indicating light-induced responses of mMeOp at all four wavelengths.

We have previously demonstrated that the blue-centered (∼420 nm λ_max_)^49^ absorption spectrum of human cone blue opsin allows for the use of blue light to activate the opsin, red wavelengths to capture signaling dynamics, enabling subcellular signaling activation, and global response imaging^6,29^. However, the above-observed broad spectral sensitivity of mMeOp makes it sensitive even to red light around 600 nm, rendering it incompatible for subcellular signaling control.

### 2.2 The mMeOp (and SqR) models

The protein-11CR interactions govern the absorption characteristics of opsins^50,51^. Indeed, as anticipated above, the λ_max_ value is inversely proportional to the vertical energy difference between the S_1_ and S_0_ states (ΔE_*S*1*−S*0_) of 11CR computed at the S_0_ room-temperature equilibrium structure (i.e., the dark state) of the opsin. To produce a blue shift in λ_max,_ one has to stabilize S_0_ and/or destabilize S_1_ (opposite changes will instead generate a red shift). The S_0_ and S_1_ energies are primarily sensitive to the opsin electrostatics (i.e., charge distributions of the opsin residues surrounding 11CR) and conjugated backbone conformation (i.e., twisting of single and double-bonds along the 11CR conjugated chain). The electrostatic modulation is possible because S_0_ and S_1_ have electronic structures leading to distinct charge distributions. More precisely, S_0_ features a positive charge localized on the -C=NH-Schiff base linkage of 11CR, while S_1_ features a positive charge delocalized away from the -C=NH-moiety (i.e., towards the chromophore β-ionone ring). The conformational modulation is instead explained by the modification of the electronic structure imposed by the twisting about single bonds causing, effectively, a decreased conjugation leading to a blue-shift or twisting about double-bonds causing a weakening of the π-bond leading to a red-shifting^21^.

An *in silico* model of WT mMeOp is essential when searching for mutations to spectrally blue-shift mMeOp to resist activation by red light. In the absence of WT structural data, we exploited the ∼40% sequence similarity between SqRh (PDB ID 2Z73) and mMeOp to produce a homology model using the MODELLER program^52^. Among an ensemble of ∼thousand models generated with MODELLER, the homology model with the highest 3D profile and quality scores was selected, refined by adjusting side-chain torsion angles, and optimized by CHARMM force field^53^, utilizing the implicit generalized Born membrane/water model^52,54^.

Fig. 2a shows the crystallographic structure of SqRh and the optimized homology model for WT mMeOp. The data show that the amino acids forming the 13C hosting cavity were conserved in the mMeOp model. In SqRh, the H-bond network between E180, Y277, and S187 residues holds the counterion E180 close to the Schiff base nitrogen (SBN), stabilizing the ground state. Furthermore, N87 and Y111 stabilize the ground state since their dipoles are oriented toward the SBN (Fig. 2b). The cavity of mMeOp exhibits an H-bond network between S167, Y255, and E160, conserved from SqRh, and it extends to a molecule of water (Wat-1), which does not affect the conformational relaxation of the counterion. As a consequence, it could be hypothesized that the negatively charged E160 cooperates with the dipole moment of Wat-1 in electrostatically stabilizing the mMeOp S_0_ with respect to S_1_ (i.e., the positive charge on the Schiff base moiety of 11CR in S_0_). Also, one can hypothesize that S_0_ is stabilized by Wat-2, that is H-bonded to the backbone carbonyl of Y91 (Fig. 2b).

**Figure 2.**
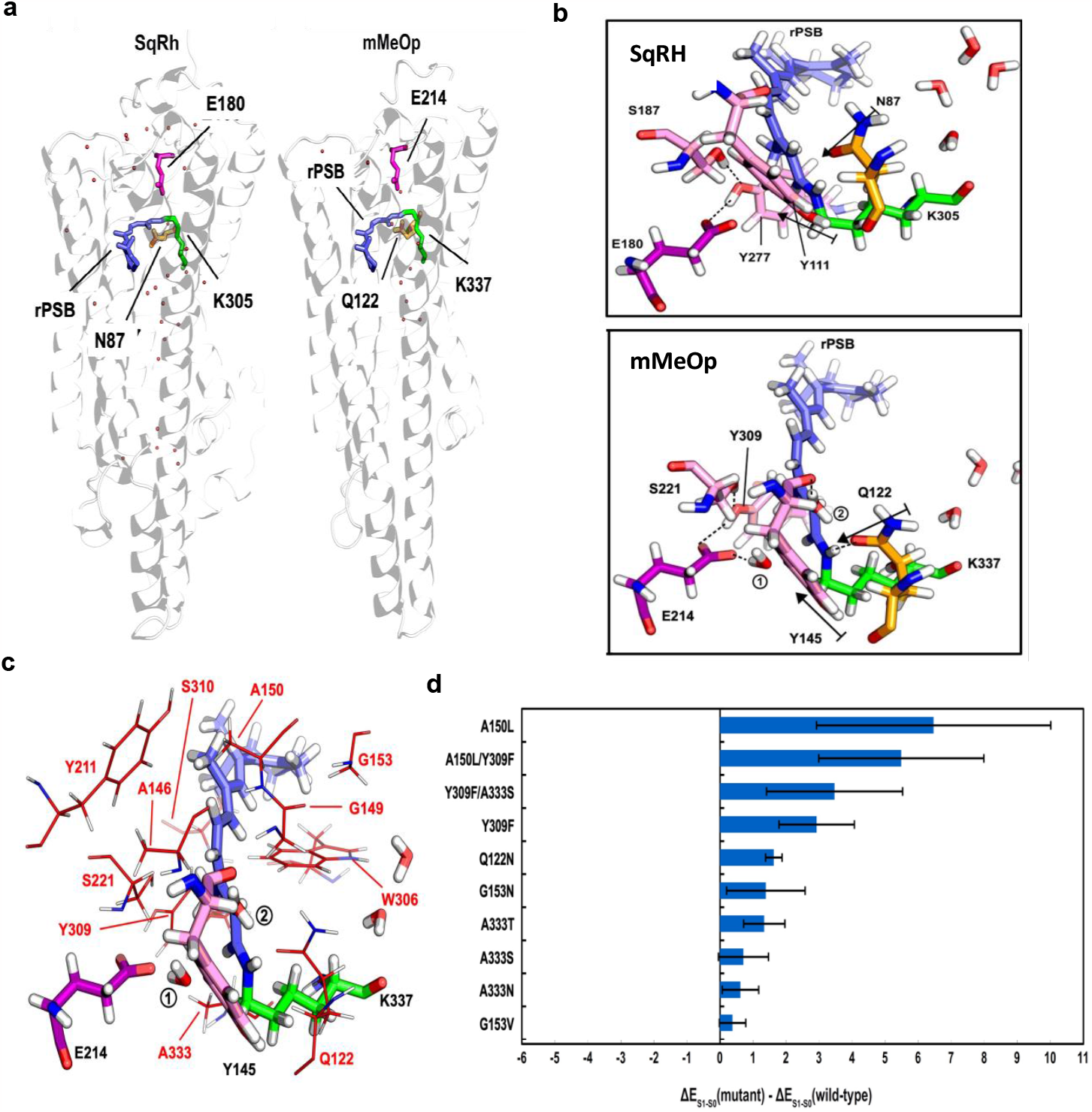
**(a)** Crystallographic structure of Squid Rhodopsin (PDB ID 2Z73) and the selected homology model of mMeOP. Retinal chromophores are shown in blue color, retinal binding sites (K305 in SqRh and K337 in mMeOp) are shown in green, main counterions (E180 and E214) are shown in purple, secondary counterions (N87 and Q122) are shown in orange, and red dots indicate water molecules. **(b)** Retinal binding cavities of QM/MM ARM models for SqRh and mMeOp. The retinal and the Schiff-base forming lysine are shown in blue and green, respectively. The primary and second counterions are shown in purple and orange, respectively. The rest of stabilizing elements are shown in pink. Labels 1 and 2 correspond with the number of water molecules in the active site of MeOp. The dashed lines and black arrows correspond to H-bonds and dipole moments, respectively. **(c)** Cavity residues of ARM model of mMeOp. Selected amino acids for point mutations are shown in red lines. The retinal chromophore is shown in blue, lysine linker (K337) in green, and the primary main counterion (E214) in purple. Water molecules are shown in red and white. **(d)** Effect of mutations on the λmax of mMeOp, computed as the difference in ΔE(S_1_−S_0_) between mutant and WT models. Blue bars correspond to the blue-shift in the absorption maximum. Black lines coincide with the standard deviation.

To computationally analyze the spectral features of mMeOp, we used the selected homology model as the input to the ARM protocol^54^. ARM generates ten QM/MM models that are then used to predict the average ΔE_S1−S0_ and, therefore, the λ_max_ value. In Table S1, we show that the ARM-generated QM/MM models can satisfactorily reproduce the experimental quantity (or, better, the difference between the λ_max_ of mMeOp and SqRh).

Since the constructed mMeOp and SqRh QM/MM models reproduce the observed λ_max_ change, we used them to investigate the mechanism responsible for the λ_max_ difference between the two opsins. Here, we limit ourselves to analyzing the λ_max_ difference in terms of (i) electrostatic interactions between the cavity and the retinal chromophore and (ii) the effect of the cavity on the chromophore conformation. To disentangle electrostatic and conformational effects, we compared the energy computed for the retinal chromophore in its protein environment (ΔE_S1−S0_ Protein Environment) and *in vacuo* (ΔE_S1−S0_ Vacuo) taken with its cavity distorted conformation. This analysis (see Table S2) revealed that the isolated 11CR taken with its cavity-distorted conformation has an ΔE_S1−S0_ of about 10-15 kcal/mol lower than inside the opsin cavity, translating into a strongly red-shifted λ_max_. In other words, the mMeOp cavity-induced 11CR geometrical changes cause a large decrease in the ΔE_S1−S0_ values. Thus, the protein electrostatics (see Table S2) must induce a large blue shift in both models, demonstrating that opposite conformational (red-shifting) and electrostatic (blue-shifting) cavity effects are responsible for the observed λ_max_ values.

To rationalize the mMeOp and SqRh λ_max_ difference, we compared the torsional distortions along the two chromophores. It is evident from Table S2 that the conformational contribution alone (**Δ**E_S1−S0_ Vacuo) would lead to mMeOp with a blue-shifted λ_max_ with respect to SqRh, contrary to the computed ΔE_S1−S0_ value for the full protein model. This result points to a dominating electrostatic effect that would blue-shift SqRh more than mMeOp. Notice that the QM/MM models of SqRh and mMeOp show C10-C11 and C12-C13 single bonds twisted more than ∼10^°^ (Fig. 3c), consistently with a backbone featuring a strong counterclockwise helicity and a blue-shifted λ_max_. However, in addition, both chromophores have a C11-C12 double bond that is ∼10^°^ twisted, leading to a dominating red-shifting distortion (Fig. 3c).

**Figure 3.**
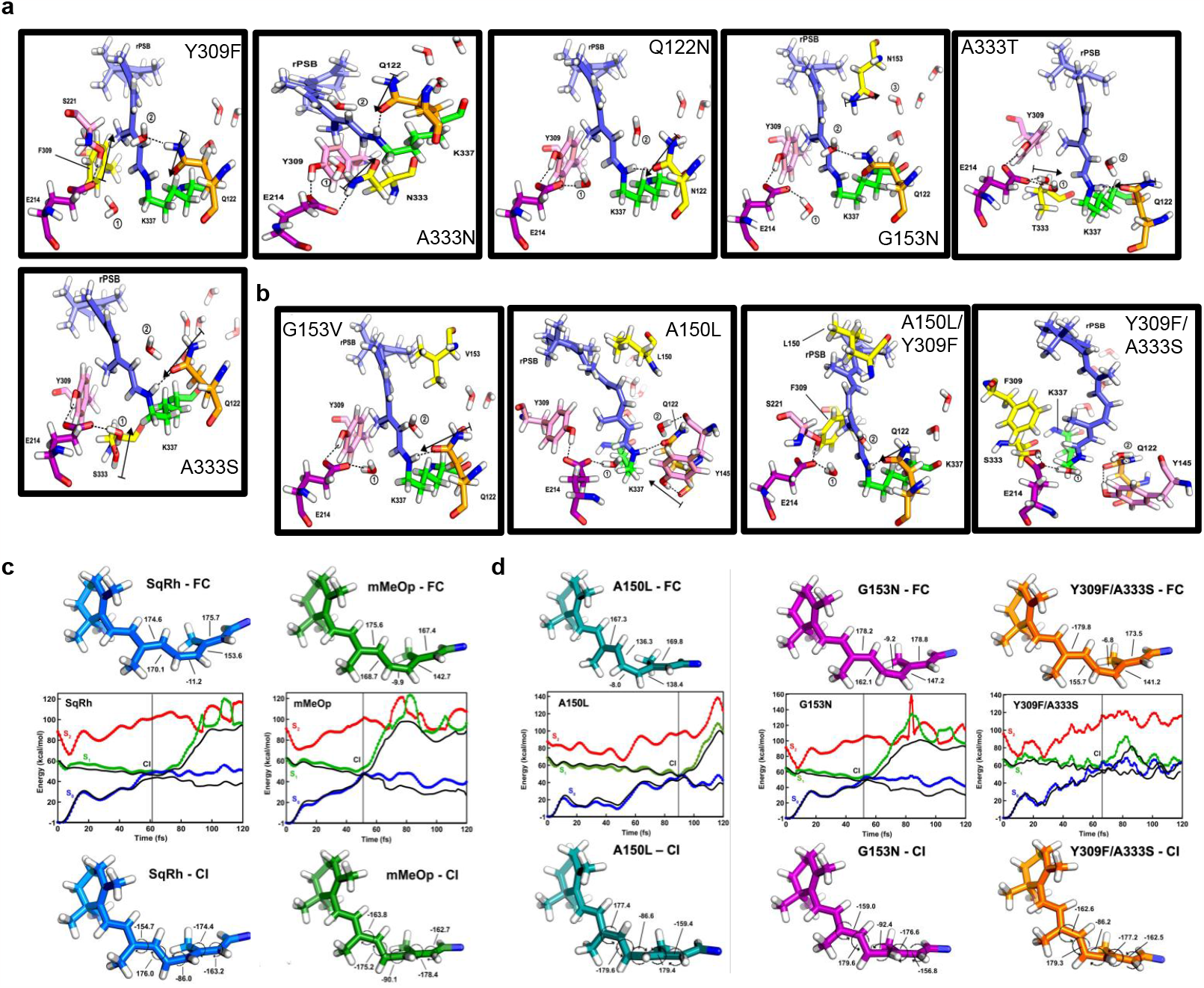
**(a)** Cavities of the blue-shifted mutants, where mutated residues (yellow) modulate ΔE(S_1_−S_0_) via electrostatic interactions, stabilizing the ground state (all except G153N). The cavity of a blue-shifted mutant, where the mutated residue (yellow) modulates ΔE(S_1_−S_0_) via electrostatic interactions, destabilizing the first excited state (G153N). The black dotted lines and black arrows indicate H-bonds and dipoles, respectively. **(b)** Cavities of the blue-shifted mutants, G153V and A150L, where mutated residues (yellow) aim to modulate ΔE(S_1_−S_0_) via steric interactions. Cavities of the blue-shifted mutants, Y309F/A333S and A150L/Y309F, where two residues are mutated (yellow) to modulate ΔE(S_1_−S_0_) both via steric and electrostatic interactions. The black dotted lines and black arrows indicate H-bonds and dipoles, respectively. **(c)** QM/MM trajectories of SqRh and mMeOp-WT ARM models, computed at the two root SA-scaled CASSCF(12,12)/6-31G*/Amber (black lines) level of theory and corrected at 3 root CASPT2 level. S_0_ (blue lines), S_1_ (green lines), and S_2_ (red lines). Franck-Condon geometries are at the top, and conical intersection geometries are at the bottom for each studied opsin. Dihedral angles are given in degrees. **(d)** QM/MM trajectories of G153N and A150L mMeOp mutants’ ARM models, computed at the two root SA-scaled CASSCF(12,12)/6-31G*/Amber (black lines) level of theory and corrected at 3 root CASPT2 level. S_0_ (blue lines), S_1_ (green lines), and S_2_ (red lines). Franck-Condon geometries are given at the top, and conical intersection geometries are provided at the bottom for each studied opsin. Dihedral angles are given in degrees.

Animal opsins usually undergo ultrafast (i.e., sub-picosecond) isomerization characterized by ultrashort S_1_ lifetimes. This has been measured for Bovine and Octopus rhodopsins with an S_1_ lifetime of less than 100 fs^55^. It has also been computationally found that both SqRh and human melanopsin (hMeOp) undergo S_1_ to S_0_ decay within 100 fs^56^. More in general, blue-shifted rhodopsins are associated with shorter S_1_ lifetimes due to the increased kinetic energy gained during S_1_ torsional relaxation toward the S_1_/S_0_ conical intersection (CoIn) funnel^40,57^, delivering the chromophore to S_0_. On the contrary, proteins with red-shifted absorption values are expected to exhibit a longer S_1_ lifetime^40^. Before using the mMeOp model for further (*e*.*g*., mutational) studies, it is therefore important to examine whether it consistently reproduces the λ_max_ value and if the model predicts a short, excited state lifetime. To do so, we performed deterministic quantum-classical Franck-Condon (FC) trajectories, namely surface-hop trajectories released on the S_1_ potential energy surface starting from the S_0_ equilibrium structure with zero initial velocities. These calculations are used to qualitatively estimate the S_1_ lifetime. Since we are only interested in the opsin evolution on S_1_, the trajectories were propagated until entering the region of an S_1_/S_0_ CoIn where decay to S_0_ occurs and formation of the ATR photoproduct is initiated. FC trajectories of SqRh and mMeOp WT models are shown in Fig. 3c. According to which both models decay within ∼70 fs; however, mMeOp reaches the hop point ∼10 fs earlier than SqRh: a result consistent with previous studies on hMeOp^56^. Even though a blue-shifted λ_max_ suggests a larger S_1_ destabilization and, therefore, a shorter S_1_ lifetime (i.e., as reported in Table S1, the ΔE_S1−S0_ value of the mMeOp WT model is larger than that of the SqRh model and, thus, an increased S_1_ relaxation and photoisomerization speed in the mMeOp models are expected^40,58^, this can also be attributed to the differences in the chromophore geometries at the FC and CoIn points. The mMeOp retinal geometry shows the lowest RMSD value (0.13966) between its FC and CoIn geometries and an average S_1_-S_2_ gap value similar to SqRh (the S_1_-S_2_ gap may contribute to slowing the dynamics down)^59^. Therefore, mMeOp would need less time to reach the CoIn than SqRh. Below, we show how we have conveniently used the QM/MM model of WT mMeOp discussed above to select, *in silico*, blue-shifting mutations expected to enhance and decrease the sensitivity to red light and isomerization timescale simultaneously.

### 2.3 Modeling, cavity analysis, and trajectory analysis of potentially blue-shifted mMeOp mutants

Enhancing the sensitivity of the opsin to blue light requires mutations that alter the interaction of the chromophore with the surrounding environment^39,40,54,60^. We explored *in silico* mMeOp mutations designed to modulate both the steric and electrostatic effects of the cavity on 11CR. Using the ARM protocol and the WT mMeOp homology model as a template, we produced the QM/MM models of eight single-residue and two double-residue blue-shifted mutants from a pool of mutations that were supposed to either stabilize S_0_ or destabilize S_1_. As displayed in Table S3, the QM/MM models of the selected mutants predicted a 5 -42 nm blue shift in λ_max_ with respect to the QM/MM WT model. Fig. 2c shows the potential mutation sites and water molecules, and Fig. 2d shows the corresponding calculated ΔE_S1−S0_ for each mutant model.

In order to examine the steric and electrostatic effects associated with each mutation, we re-computed their λ_max_ after zeroing the charges of the mutated residues. It was found that Y309F, Q122N, A333T, A333S, and Y309F/A333S mutants only slightly affected the retinal chromophore conformation (see Fig. 3a and 3b) and that it is the change in side-chain polarity that increases the S_1_-S_0_ energy gap (Fig. S1). In contrast, the A333N and G153N mutants display a blue-shifting effect caused by sterically induced changes in the 11CR conformation. In fact, for instance, the N153 residue largely twists the single bonds of 11CR (Fig. 3a) and consequently, the computed λ_max_ value decreases with respect to the WT (Fig. S2).

In an attempt to increase such a blue-shifting effect, we additionally mutated G153 into a *Val* (G153V) in G153N. However, the effect of such mutation on the ΔE_S1−S0_ was less than expected (Fig. S2). Finally, the same calculations showed that models A150L and A150L/Y309F produced the largest blue-shifting. While Fig. S2 indicated that the inserted bulkier side chains twist the 11CR conjugated chain into a red-shifted configuration, the electrostatic effect of the cavity must produce a dominating blue-shift. The modeled retinal binding cavities of G153V, A150L, Y309F/A333S, and A150L/Y309F are depicted in Fig. 3b. To qualitatively estimate the S_1_ lifetime of the mMeOp mutants and ensure that it is consistent with a functional opsin (i.e., it is similar to the lifetime of the WT model), we performed FC quantum-classical trajectories. In the past, short S_1_ lifetimes were assumed to indicate the sensitivity (i.e., isomerization quantum yield) of mMeOp to light as indicated by the Landau-Zener model relating faster isomerization with higher sensitivity^61^. However, it has been recently demonstrated, by computing entire sets of quantum-classical trajectories representing the opsin’s S_1_ population dynamics, that while vertical excitation energies and excited state lifetimes are inversely proportional, excited state lifetimes and isomerization quantum yields are not proportional^56^. This was demonstrated by comparing SqRh with hMeOp and depends on the fact that the Landau-Zener model must be applied to the reaction coordinate locally, driving the decay event^62^.

FC trajectory calculations were performed for three promising blue-shift mutant models: G153N, A150L, and Y309F/A333S. None of the mutants displayed a decreased S_1_ lifetime. However, the lifetime remains of the same magnitude as the WT. The G153N mutant reaches the CoIn at ∼53 fs, while the A150L and Y309F/A333S mutants enter the CoIn region after ∼90 fs and ∼66 fs, respectively (Fig. 3d). As stressed above, these results are not quantitative and cannot be used to predict the “observable” lifetime. Such a prediction would only be possible after running hundreds of quantum-classical trajectories starting from a corresponding number of initial conditions. Only by performing such statistics may one expect that a reduced S_1_ lifetime is associated with an increased S_1_-S_0_ gap^40^. Such a demanding calculation goes beyond the scope of the present research. In the specific case of the computed A150L FC trajectories, the analysis of retinal geometries and energies indicates that the S_1_ lifetime for each mutant is consistent with RMSD analysis and CASPT2 calculations (Fig. 3d), pointing to an isomerization that would slow down the reaction. However, structural considerations cannot explain why the G153N and Y309F/A333S mutants show a speed slower than expected. In this case, the computation of CASPT2 energies revealed a ∼4 kcal/mol S_1_-S_2_ gap, which strongly increases the possibility of interaction between these two states, leading to a slower isomerization mechanism^59^. Again, it must also be pointed out that the above considerations are qualitative since they are based on a single FC trajectory. A statistical analysis sampling different initial conditions could substantially refine the presented relationship between excitation energies and excited state lifetime/reaction speed.

### 2.4 Spectral sensitivity and signaling of the predicted blue-shifted mMeOp mutants

Using site-directed mutagenesis, we generated the ten *in silico*-predicted blue-shifted MeOp mutants (Fig 2d). Next, using live cell imaging, we examined whether the mutants were sufficiently blue-shifted to resist activation by red light (594 nm) and whether they possessed signaling activities similar to the WT MeOp exposed to wavelengths at or below 500 nm. We first tested Gq signaling by expressing each mMeOp mutant using mCh-PH in HeLa cells, and mutants that did not show PIP2 hydrolysis were selected for subsequent screening. We imaged cells using 594 nm red light (1.5 μW). Except for Y309F, Q122N, A333N, and A333S, which showed PIP2 hydrolysis upon red light exposure (Fig. S3 a-d), all other mutants (A150L, G153N, G153V, A333T, Y309F/A150L, and Y309F/A333S) did not respond to red light activation (Fig. 4a and Fig S3 e, f). We next examined whether the red-light-resisting mutants could sense blue light and activate signaling. Similar to Fig. 1c, we expressed each of the six mMeOp mutants alongside mTq-PH in HeLa cells. Although G153V and Y309F/A150L expressing cells did not show PIP2 hydrolysis when exposed to 445 nm, 1.5 μW, blue light (Fig. S3 e and f), indicating a complete loss of activity. A333T, A150L, G153N, and Y309F/A333S expressing cells showed robust PIP2 hydrolysis upon blue light (Fig. 4b).

**Figure 4.**
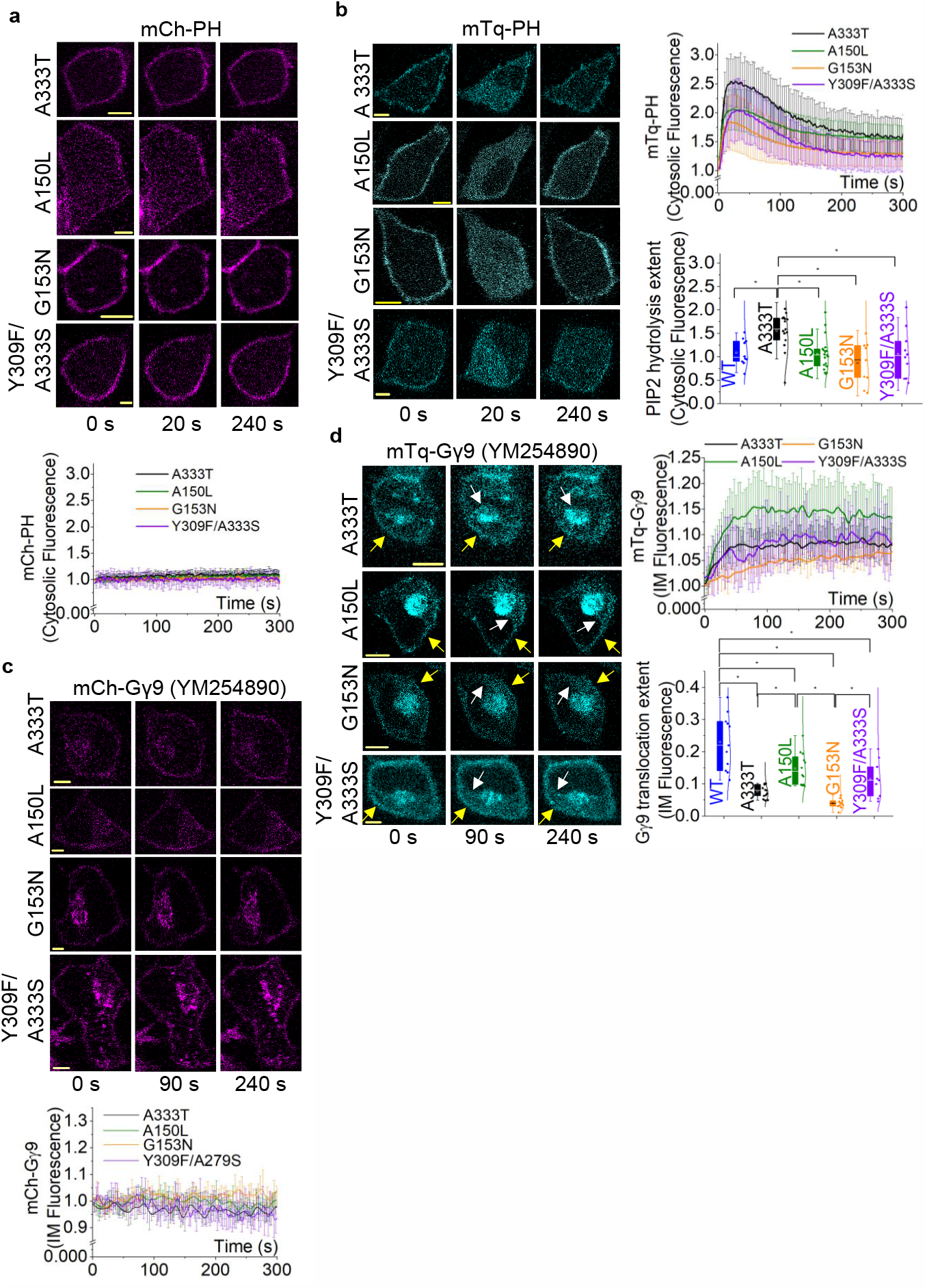
MeOp blue-shift mutants exhibit robust Gq and Gi/o activity in blue light but resist red light-induced activation. **(a)** Red light did not induce detectable hydrolysis of PIP2 with any of the mutants. A333T (n = 11), A150L (n =11), G153N (n = 8), and Y309F/A279S (n = 11). **(b)** HeLa cells expressing mTq-PH alongside MeOp-blue-shift mutants; A333T (n = 15), A150L (n = 19), G153N (n = 9), Y309F/A279S (n = 10) exhibit robust PIP2 hydrolysis upon photoactivation by blue light in the presence of 10 μM 11CR. Images and the corresponding plot show the PIP2 sensor dynamics in the cytosol. The whisker box plot shows the PIP2 hydrolysis extent of each mutant and WT. **(c)** A333T, A150L, G153N, and Y309F/A279S mutants resist photoactivation by red light (n = 12: A333T, 9: A150L, 10: G153N, 11: Y309F/A279S). **(d)** HeLa cells expressing mTq-Gγ9 alongside MeOp-blue-shift mutants; A333T, A150L, and G153N, and Y309F/A279S exhibit Gγ9 translocation from the PM to IMs upon photoactivation by blue light in the presence of 10 μM 11-*cis*-retinal and the Gq inhibitor, YM254890. White and yellow arrows indicate the fluorescence intensity increase on IMs and loss of fluorescence intensity on the PM, respectively. The corresponding plots show Gγ9 dynamics on the IMs (n = 13: A333T, 12: A150L, 11: G153N, 9: Y309F/A279S). The whisker box plot shows the Gγ9 translocation extent of each mutant and WT. Average curves were plotted using ‘n’ cells, n = number of cells. Error bars represent SD (standard deviation). Statistical comparisons were performed using One-way-ANOVA; p<0.05, (**\***: population means are significantly different). The scale bar = 5 μm.; mTq: mTurquoise; mCh: mCherry

Next, we compared the extent of PIP2 hydrolysis induced by WT with each functional mutant. MeOp-A333T induced a significantly higher extent, while A150L, G153N, and Y309F/A333S mutants showed similar PIP2 hydrolysis properties to the WT (Fig. 4b, whisker plot, One-way ANOVA: F_4,62_ = 5.35772, p = 9.67E-4, Table S4 a and b).

Since WT mMeOp activates both Gq and Gi/o pathways efficiently^63^, we tested whether these mutants also induce efficient Gi/o signaling upon blue light in the presence of YM-254890. All four mutants showed robust Gγ9 translocation upon blue light exposure (Fig. 4d). However, compared to the WT, Gγ9 translocation responses induced by the mutants were significantly lower (Fig. 4d, whisker plot, One-way ANOVA: F_4,55_ = 22.38526, p = 1.02508E-10, Table S5 a and b), indicating a reduced Gi/o activity, likely due to the mutagenesis. In a control experiment, we observed that these mutants did not show Gγ9 translocation when exposed to red light (Fig. 4c). Additionally, G153N mutant exhibited the lowest Gq and Gi/o activity of all, indicating the mutation in the retinal binding cavity has compromised its overall activity (Fig. 4b and d, whisker plots).

### 2.5 Spectral characterization of red light-resisting mMeOp mutants resisting red light activation

To determine the absorption spectra, we expressed and purified the mMeOp mutants (A333T, A150L, G153N, and Y309F/A333S) in HEK293S cells. The absorption spectrum for mMeOp A333T yielded a λ_max_ of ∼450 nm (∼20 nm blue-shifted with respect to the WT (Fig. 5a). Since it was not possible to determine the spectral blue shift of other mutants using absorption spectroscopy (likely due to low expression levels or low stability in detergent), we determined their action spectra instead by measuring Gq pathway activation-induced Ca^2+^ mobilization upon activation of MeOp-WT and mutants at wavelengths ranging from UV to red using the aequorin reporter assay^64,65^. In the presence of 5 μM Coelentrazine-H substrate and 10 μM 11CR, the baseline luminescence readings were collected from HeLa cells expressing mMeOps (WT, A333T, A150L, G153N, or Y309F/A333S) and aequorin. Afterward, each well was exposed to near monochromatic light of the selected wavelength while collecting aequorin luminescence (Fig. 5c). We captured luminescence data at ∼20 different wavelengths spanning the 230-700 nm range.

**Figure 5.**
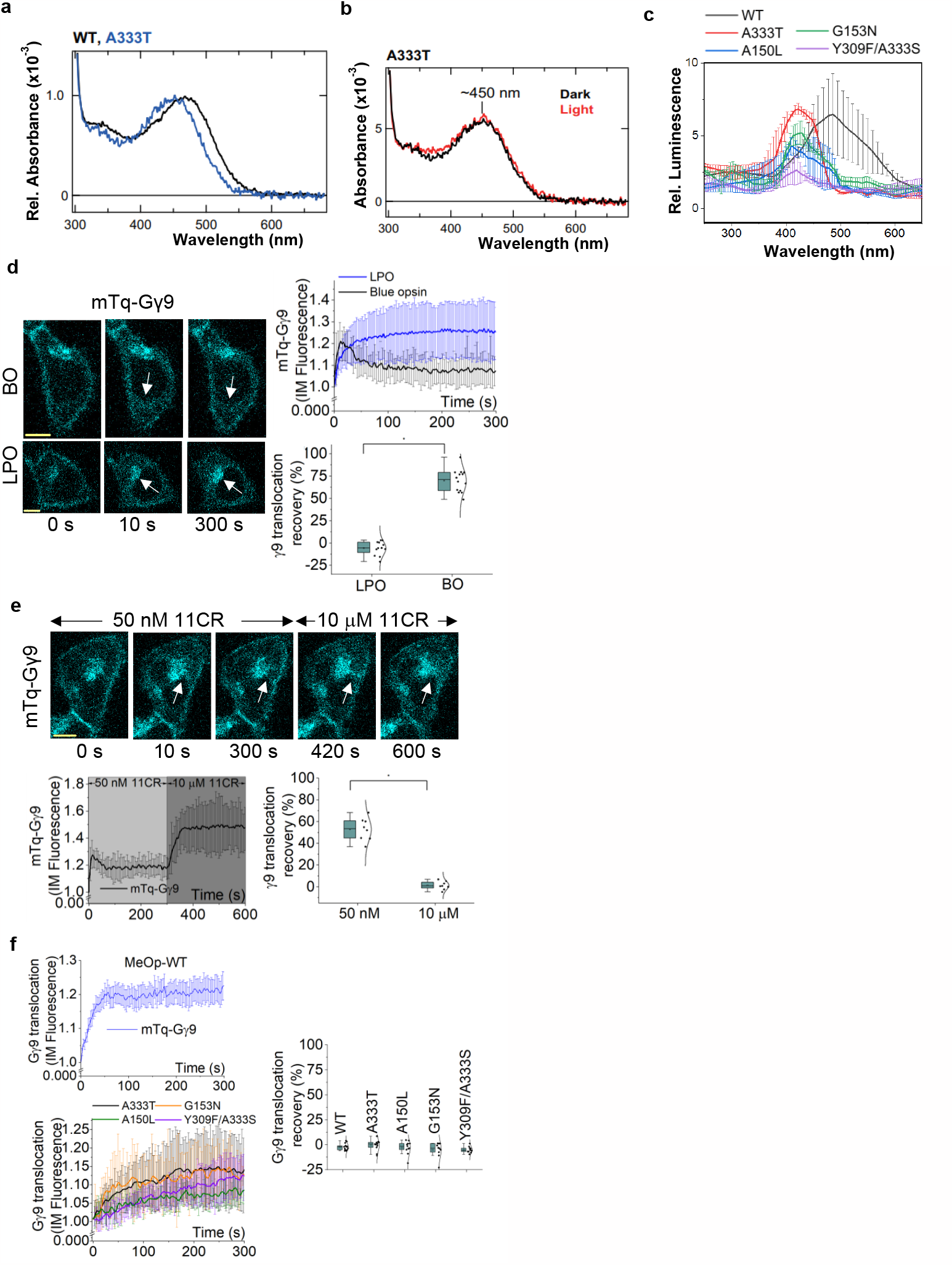
Experimental absorption and action spectra of MeOp mutants show significant blue shifts compared to the WT and are bistable. **(a)** Comparison of the WT and A333T mutant MeOps’ absorption spectra show a ∼20 nm blue shift in A333T. The blue and black plots indicate the absorption spectra of MeOp-A333T and WT-MeOp, respectively. (**b)** Purified MeOp-A333T mutant from heterogeneously expressed HEK293S cells exhibited a λ_max_ of ∼450 nm in the dark (black plot). The light-exposed spectrum of MeOp-A333T was nearly identical to the dark spectrum with a similar λ_max_ (red plot). **(c)** The action spectra of A333T, A150L, G153N, and Y309F/A333S-MeOp exhibit significant blue shifts compared to the WT. The action spectrum of WT-MeOp (black) exhibits an action maximum (λ^a^_max_) of ∼486 nm The action spectrum of MeOp-A333T (red) exhibited a λ^a^_max_ of ∼421 nm, while λ^a^_max_ of A150L (blue), G153N (green), and Y309F/A333S (purple) were ∼411, ∼419 nm, and ∼420 nm, respectively. **(d)** HeLa cells expressing blue opsin and mTq-Gγ9 exhibit rapidly attenuating Gγ9 translocation upon blue opsin activation in the presence of 50 nM 11CR (n = 14). In contrast, cells show sustained Gγ9 translocation upon blue light activation of lamprey parapinopsin with 50 nM 11CR (n = 12). Grey arrows indicate the fluorescence intensity changes in the IMs due to Gγ9 translocation and attenuation. The whisker box plot shows the recovery extent of translocated Gγ9. **(e)** The attenuated Gγ9 translocation response with blue opsin in low retinal concentration (50 nM 11CR) can be rescued by increasing the retinal concentration up to 10 μM. Grey arrows indicate the fluorescence intensity changes in the IMs. The corresponding line plot shows the mTq-Gγ9 dynamics in the IMs. The whisker box plot shows the attenuation extent of translocated Gγ9 with the two retinal concentrations. (n = 8) **(f)** The bistable, MeOp-WT exhibits non-attenuating, sustained Gγ9 translocation with blue light activation in the presence of 50 nM 11CR. Similarly, the mutants, A333T, A150L, G153N, and Y309F/A333S exhibit sustained Gγ9 translocation upon blue light activation with 50 nM 11CR. The line plots show the mTq-Gγ9 dynamics in the IMs. The whisker box plot shows the recovery extent of translocated Gγ9 of WT (n = 13), A333T (n = 11), A150L (n = 11), G153N (n = 10), and Y309F/A333S (n = 10). Average curves were plotted using ‘n’ cells, n = number of cells. Error bars represent SD (standard deviation). Statistical comparisons were performed using one-way-ANOVA; p<0.05, (**\***: population means are significantly different).

We observed wavelength-dependent aequorin luminescence increase for the WT and mutant MeOps. The corresponding luminescence increases were calculated and fitted (Fig. 5c). Action spectra of WT mMeOp showed an action spectrum maximum (λ^a^_max_) of 486 nm (Fig. 5c), a relatively close value to the 470 nm λ_max_ obtained from its absorption spectrum, validating the feasibility of using action spectra to examine the spectral characteristics of opsins^64^. As expected, based on the signaling data, the A333T mutant showed a λ^a^_max_ at 421 nm, a 65 nm blue-shift compared to the WT (Fig. 5c) The differences between absorption spectra in Fig 5a and the corresponding action spectra in Fig. 5c can be partially due to the protein environment differences between detergent and living cells. Action spectra of the A150L, G153N, and Y309F/A333S mutants showed λ^a^_max_ of 411 nm (75 nm blue-shift), 419 nm (67 nm blue-shift), and 420 nm (66 nm blue-shift), respectively (see Table S3 and Fig. 5c).

### 2.6 Blue-shift melanopsin mutants are bistable

Since engineering subcellular signaling-capable optogenetic opsins is a key goal of this work, we examined whether the four selected blue-shifted mMeOp mutants also possess bistability similar to the WT. When irradiated, bistable opsins in their dark-adapted state (i.e., kept in the dark until fully equilibrated), a stable photoproduct called light-adapted state is generated, resulting in a photo-product with a similar or slightly altered absorption spectra^66^. It has been shown that WT mMeOp forms a slightly red-shifted light-adapted state^28^. Due to the above reasons, we could not obtain resolved light-irradiated spectra for A150L, G153N, and Y309F/A333S. However, the light-irradiated spectrum of the A333T mutant showed a similar spectrum to that of the dark-adapted state (Fig. 5b, red plot). This indicated that not only A333T mutant is bistable, but other mutants are also likely to be bistable. Next, we indirectly tested mutants’ bistability using their ability to recycle retinal, a characteristic feature of most bistable opsins, including mMeOp. We hypothesized that at a low enough 11CR concentration, a monostable opsin should show only a transient signaling activity, while the bistable opsins will show sustained signaling.

We used well-characterized monostable human cone blue opsin (BO) and bistable lamprey parapinopsin (LPO) expressing cells to validate this assay. In the presence of a relatively lower concentration of 11CR (50 nM), BO-expressing cells showed a robust, however, transient mTq-Gγ9 translocation from the plasma membrane to endomembranes upon exposure to blue light (Fig. 5d, BO: image of 10 s and plot). On the contrary, under the same 11CR availability, bistable LPO showed a sustained Gγ9 translocation (Fig. 5d, LPO: image of 10 s, and plot). We calculated the attenuation/recovery percentage of the Gγ9 translocation for both BO and LPO in the presence of blue light with 50 nM 11CR. Although the Gγ9 translocation attenuation was negligible with LPO, BO-induced Gγ9 translocation showed a significantly higher attenuation of 67.8 ± 3.0 % (Fig. 5d, whisker plot, One-way ANOVA: F_1,23_= 384.92321, p = 7.35E-16, Table S6 a and b).

As a control, when we added 10 μM 11CR to a Gγ9 translocation-attenuated cell in the presence of 50 nM 11CR (Fig. 5e, images of 10 s and 300 s), Gγ9 showed re-translocation (Fig. 5e, images at 420 s and 600 s). The Gγ9 translocation attenuation percentage with 50 nM 11CR was significantly higher than that with 10 μM 11CR (Fig. 5e, whisker plot, One-way ANOVA: F_1,15_ = 174.93367, p = 2.66E-9, Table S7 a and b). These observations indicated that this assay could distinguish functional differences between monostable and bistable opsins. Nevertheless, this assay should be cautiously used, considering each of the stable state’s signaling propensities and spectral properties.

Using this strategy, we next examined the bistable nature of the mMeOp WT and mutants: A333T, A150L, G153N, and Y309F/A333S in HeLa cells also expressing with mTq-Gγ9. When we exposed cells to blue light in the presence of 50 nM 11CR, WT mMeOp induced sustained Gγ9 translocation, further indicating the validity of this assay (Fig. 5f). A333T, A150L, G153N, and Y309F/A333S mutants also induced non-attenuating Gγ9 translocation responses upon blue light exposure (Fig. 5f). Like LPO, the calculated % recovery of Gγ9 translocation was negligible (Fig. 5f, whisker plot). This data indicated that the above four blue-shifted MeOp mutants have retinal-recycling capabilities, suggesting they are bistable.

### 2.7 A333T MeOp mutant allows spatio-temporal control of single cells and subcellular G protein signaling

Since our data indicate that the A333T mutant and WT mMeOp have nearly similar Gq signaling activities (Fig. 4a), considering its newly gained red light resistivity, we examined the feasibility of using A333T MeOp for temporal signaling control while continuously imaging cells using red light. Unlike the WT (Fig. 6a, images, and plot), A333T expressing cells did not show PIP2 hydrolysis upon imaging mCh-PH (Fig. 6b, images up to 55 s, and plot’s magenta background). At 60 s, we exposed cells to both red and blue light. Cells showed robust PIP2 hydrolysis (Fig. 6b, images of 90 s, and plot’s blue background). This indicated the feasibility of using A333T mutant to capture pre-and post-stimulation signaling and cell behavior using red light with a user-defined temporal control.

**Figure 6.**
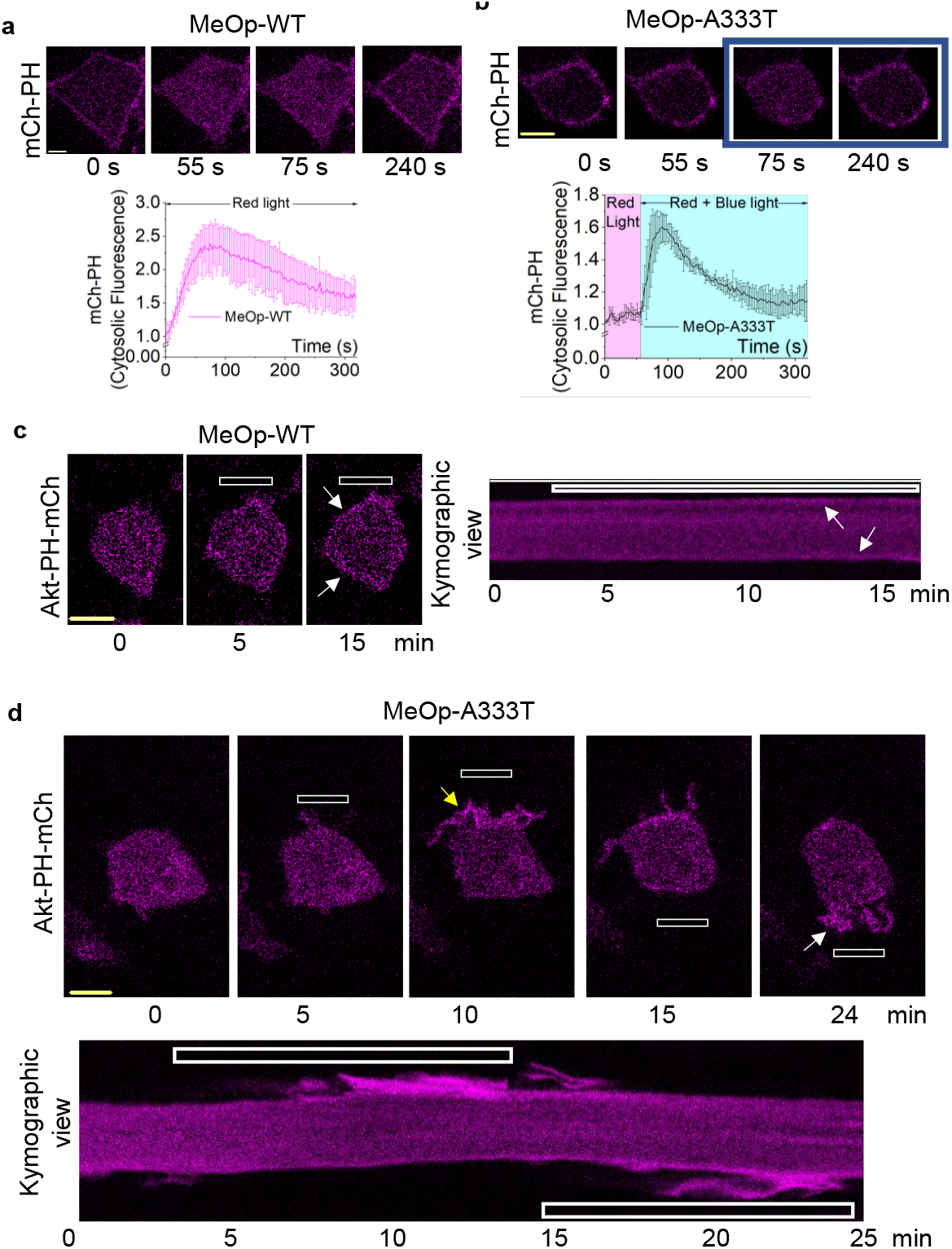
MeOp-A333T mutant allows for spatio-temporal and subcellular G protein signaling control in living cells. **(a)** HeLa cells expressing MeOp-WT and mCh-PH exhibit robust PIP2 hydrolysis upon red light illumination in the presence of 10 μM 11CR. The corresponding plot shows mCh-PH dynamics in the cytosol. (n = 12) **(b)** MeOp-A333T-expressing cells do not show a detectable PIP2 hydrolysis with red light, however, exhibited robust PIP2 hydrolysis upon blue light exposure. The corresponding plot shows mCh-PH dynamics in the cytosol with red light (magenta background) and blue light (blue background) (n = 10). **(c)** RAW264.7 cells expressing MeOp-WT and Akt-PH-mCh exhibit global PIP3 (white arrows) due to red light imaging despite localized blue light. The white box indicates the confined region of blue light. Although data is shown only from one cell, experiments were conducted in multiple cells to test the reproducibility. **(d)** RAW264.7 cells expressing MeOp-A333T and Akt-PH-mCh exhibit localized PIP3 upon localized blue light activation (yellow arrows). Upon changing the blue light stimulus to the other side of the cell, PIP3 is generated from the blue light-exposed side (white arrows). The kymographic view of the cell indicates localized PIP3 generation with locally confined blue light. Although data is shown only from one cell, experiments were conducted in multiple cells to test the reproducibility. Average curves were plotted using ‘n’ cells, n = number of cells. Error bars represent SD (standard deviation).

Previously, we have shown localized blue opsin activation and the resultant Gβγ-induced localized PIP3 and directional migration in live cells^30,67^. However, unlike RAW264.7 macrophages, due to the lack of appropriate Gγ types, HeLa cells do not show PIP3 generation upon Gi/o GPCR activation^46^. Therefore, to examine MeOp-induced localized PIP3 generation, we used RAW264.7 cells. RNAseq data shows that the endogenous Gαq level in RAW264.7 cells is significantly lower than other Gα subtypes^67^. Therefore, as described previously^67,68^, to increase Gβγ contribution from the Gq pathway, in addition to A333T or WT-MeOP and Akt-PH-mCh (PIP3 sensor), we also expressed Gαq-CFP in RAW264.7 cells. Next, we exposed one side of the cell to blue light (Fig. 6c, image of 10 min: white box) while imaging Akt-PH-mCh using red light. Imaging WT-mMeOp expressing cells using red light activated the opsin globally, inducing global PIP3 generation, which we did not observe for the mutant (Fig. 6c, white arrows, kymograph, and movie S1). The localized blue light here failed to induce subcellular signaling activation. Interestingly, MeOp-A333T activation induced localized PIP3 generation and migration of the cell towards the blue light (Fig. 6d, image of 10 min, yellow arrow, kymograph, and movie S2). Upon switching the blue light stimulus to the opposite side, the PIP3 generation was also switched (Fig. 6d, image of 24 min, white arrow, kymograph, and movie S2). This data demonstrates that similar to blue opsin, the spectrally blue-shifted MeOp variants, especially A333T-MeOp, can be used for spatiotemporal control of single-cell and subcellular signaling.

## Discussion

We presented the results of a combined computational and experimental effort to engineer a novel bistable, opsin-based optogenetic tool capable of being activated by wavelengths in the blue region but not by red light. The ability of SqRh and mMeOp QM/MM models to reproduce the experimental λ_max_ values supports the use of the ARM protocol in assisting the design of blue-shifted mutants. The analysis of 11CR chromophore binding cavities of the models then informs on the roles of residue charges, dipoles, and water molecules located at different positions relative to the 11CR conjugated chain for determining the opsin λ_max_ and thus the rational design of blue-shifted mutants. Furthermore, comparing the opsin environment and *in vacuo* calculations disentangle the 11CR chromophore conformational and electrostatic effects underlying the λ_max_ change.

ARM was used to select ten blue-shifting mutant models starting from a validated WT mMeOp model. These mutants were thus constructed and investigated in the laboratory, confirming their blue-shifted absorption with respect to WT mMeOp. However, live cell imaging showed some lost functionality, likely due to unforeseen structural defects, leaving four functional mutants available for further studies (see Table S3). A computational analysis of the residue-replacement effects on ΔE_S1−S0_ variation relative to the WT as well as deterministic Franck-Condon trajectories provided mechanistic information and indicated ultrafast photo-reactivity associated with an S_1_ lifetimes <100 fs. More specifically, both WT and mutant models corroborated that the general S_1_ lifetimes are inversely proportional to the λ_max_ value. This was governed by cavity-imposed electrostatic stabilizations/destabilization and torsional distortion of 11CR. All results are consistent with a strong light sensitivity of WT mMeOp, supported by previous studies on hMeOp indicating a quantum efficiency of 54% (i.e. 54 photons over 100 lead to isomerization)^56^. The computational analysis also demonstrated that some mutants isomerize slower than WT-mMeOp, explaining why G153N and A150L, the blue-shift mutants, do not correspond to a faster isomerization rate.

Ten predicted mMeOp mutants whose models had significantly blue-shifted λ_max_, were constructed and expressed in the lab. However, as anticipated from the observed spectral blue shift in the absorption spectra, live cell imaging showed that only A333T, A150L, G153N, and Y309F/A333S resisted activation by red light; however, they did not lose functionality when exposed to blue wavelengths. Interestingly, the A333T mutant showed a significantly higher extent of PIP2 hydrolysis compared to the WT when exposed to blue light (445 nm). However, this does not essentially indicate that the A333T mutation enhanced the Gq activity of the opsin because, in this experiment, the wavelength used is much closer to the λmax of the mutant than that of the WT (both the absorption and action spectra show that A333T absorb better around 445 nm compared to the WT).

Therefore, we argue that these mMeOp mutants can be used to control subcellular signaling, which is not possible using WT MeOp. Additionally, the bistability of these mutants makes it possible to overcome a significant limitation of monostable rhodopsins, the need for continuous 11CR supply. This is an essential advantage in a versatile optogenetic tool, as (i) the photosensitization of added 11CR can cause phototoxicity^69,70^, and (ii) there is limited retinal availability in tissues outside the retina. Demonstrating the spectral selectivity, as well as subcellular signaling potential, we showed that the mMeOp A333T mutant activates localized PIP3 generation and cell migration. Considering that spatially and temporally variable stimuli likely to induce asymmetric signaling *in vivo*, GPCRs are primary drivers of physiology and pathology, for instance, in regulating cell fate driving crucial cell behaviors, including neuronal symmetry breaking, differentiation, and migration^6^, we proposed that the newly engineered, spectrally blue-shifted MeOp mutants will be beneficial optogenetic tools.

Since MeOp showed unprecedented promiscuity for both Gq and Gi/o pathways by mutating its intracellular loops, we previously engineered a Gq-selective MeOp variant^25^, allowing for Gq-exclusive, red-light-resisting hybrid MeOp engineering in the future. Availability of these opsin mutants, together with the use of implanted optrodes^71,72^ or two-photon excitation methods^73,74^, will enable optical control of the Gq pathway that influences vital physiological functions, such as Gq-induced blood pressure elevation and associated cardiac hypertrophy^75^, PI3K-induced neural plaque accumulation and inflammation in neurodegenerative diseases caused by the activation of downstream GSK3-beta and IKK a/b pathways^76^.

Further, we also help fill the need for optogenetic tools capable of controlling subcellular signaling while motoring perpetual molecular regulation underlying pathways; we believe that the presented MeOp mutants, as well as other mutants that will be prepared in the future using the proposed computational-biochemical framework, will provide dedicated opsins for subcellular and *in-vivo* applications.

## 3. Material and methods

### 3.1 Reagents

The reagents used were as follows: Q5^®^ Site-Directed Mutagenesis Kit (NEB), Gibson Assembly^®^ Master Mix (NEB), and NEB® 5-alpha Competent E. coli (NEB), 11-*cis*-retinal (National Eye Institute), YM-254890 (Focus Biomolecules), Coelentrazine-H (AAT Bioquest). Stock solutions of compounds were prepared according to manufacturers’ recommendations. Before adding to cells, all stock solutions were diluted in 1% Hank’s balanced salt solution (HBSS) or regular cell culture medium.

### 3.2 DNA constructs and cell lines

DNA constructs used were as follows: DNA constructs used for mCh-PH, GFP-PH, Venus-PH, Blue opsin-mTurquoise, and lamprey parapinopsin have been described previously^6,18,45,77,78^. Fluorescently tagged Gγ9 subunits and αq–CFP were kindly provided by Professor N. Gautam’s laboratory, Washington University, St Louis, MO. mTq-PH was made by switching the mCherry fluorescent tag in mCh-PH (in PCDNA3.1) to mTurquoise using restriction cloning. All MeOp mutants were created using site-directed mutagenesis (NEB) from the parent construct, MeOp-1D4, in the PCDNA3.1 vector. All imaging and action spectra experiments were performed using MeOp-WT and mutants in PCDNA3.1. For protein purification experiments, MeOp-WT and mutants tagged with the rho-1D4 epitope sequence (ETSQVAPA) were inserted into the pMT vector using restriction cloning. Primers were designed using Nebuilder (for Gibson assembly or restriction cloning) and NEbasechanger (for site-directed mutagenesis). Cloned cDNA constructs were confirmed by sequencing from commercial sources. Cell lines used were as follows: HeLa and RAW264.7 cells were purchased from the American Tissue Culture Collection (ATCC).

### 3.3 Cell culture and transfections

HeLa cells were cultured in minimum essential medium (Corning) supplemented with 10% heat-inactivated dialyzed fetal bovine serum (DFBS, Atlanta Biologicals) and 1% penicillin-streptomycin-amphotericin (PSA, 100X stock, Corning) and grown at 37 °C with 5% CO_2_. RAW264.7 cells were maintained in Roswell Park Memorial Institute (RPMI) 1640 medium (Corning) supplemented with 10% DFBS and 1% PSA. Cells were cultured in 35 mm, 60 mm, or 100 mm cell culture dishes (Celltreat). DNA transfections were performed using electroporation with the Cell line Nucleofector^TM^ Kit V (RAW264.7 cells) or lipofectamine 2000 transfection reagent (HeLa cells) unless otherwise specified. For electroporation, the electroporation solution was prepared with the Nucleofector solution (82 μL), Supplement solution (18 μL), and appropriate volumes of DNA constructs. For each experiment, ∼2-4 million cells were electroporated using the T020 method of the Nucleofector™ 2b device (Lonza). Immediately after electroporation, cells were mixed with cell culture medium at 37 °C and seeded onto 14 mm glass-bottomed wells (#1.5 glass coverslip) in 29 mm cell culture treated dishes. Cells were imaged ∼5-6 hours post-electroporation. Two days before imaging HeLa cells, 7 x 10^4^ cells were seeded on a 14 mm glass bottom well with a #1.5 glass coverslip in a 29 mm cell culture dish. The following day, cells were transfected with appropriate DNA combinations using the transfection reagent Lipofectamine 2000 (Invitrogen) according to the manufacturer’s protocol and then incubated in a 37 °C, 5% CO2 incubator. Cells were imaged after 16 h of the transfection.

### 3.4 Live cell imaging, image analysis, and data processing

The methods, protocols, and parameters for live-cell imaging are adapted from published work^69,79,80^. Briefly, live-cell imaging, single-cell, and subcellular photo-stimulation experiments were performed using a spinning disk Confocal Imaging System (Yokogawa CSU-X1, 5000 rpm) composed of a Nikon Ti-R/B inverted microscope with a 60X, 1.4 NA oil objective and iXon ULTRA 897BVback-illuminated deep-cooled EMCCD camera. Photoactivation and Spatio-temporally controlled light exposure on cells in regions of interest (ROI) were performed using a laser combiner with 40-100 mW solid-state lasers (445, 488, 515, and 594 nm) equipped with Andor® FRAP-PA unit (fluorescence recovery after photobleaching and photoactivation), controlled by Andor iQ 3.1 software (Andor Technologies, Belfast, United Kingdom). Fluorescent sensors such as mCh-PH, mCh-γ9, and Akt-PH-mCh, were imaged using 594 nm excitation−624 nm emission settings; Venus-PH and Venus-γ9 were imaged using 515 nm excitation and 542 nm emission; GFP-PH and GFP-γ9 were imaged using 488 nm and 510 nm emission; mTq-PH, Gαq-CFP, Blue opsin-mTq was imaged using 445 nm excitation and 478 nm emission. In experiments with MeOp, we imaged cells with the respective color to find cells with the sensor expression before adding retinal. Additional adjustments of laser power with 0.1%-1% transmittance were achieved using Acousto-optic tunable filters (AOTF). Ophir PD300-UV light meter was used for laser power measurements. Data acquisition, time-lapse image analysis, processing, and statistical analysis were performed as explained previously^69^. Briefly, Time-lapse images were analyzed using Andor iQ 3.1 software by acquiring the mean pixel fluorescence intensity changes of the entire cell or the selected area/regions of interest (ROIs). Briefly, the background intensity of images was subtracted from the intensities of the ROIs assigned to the desired areas of cells (plasma membrane, internal membranes, and cytosol) before intensity data collection from the time-lapse images. The intensity data from multiple cells were opened in Excel (Microsoft office®) and normalized to the baseline by dividing the data set by the average initial stable baseline value. Data were processed further using Origin-pro data analysis software (OriginLab®).

### 3.5 Aequorin reporter assay

HeLa cells at 75-85% confluency in a 100 mm dish were transfected with MeOp (WT or mutants) alongside aequorin with PEI transfection reagent (in DNA: PEI, 1:1 mass ratio). On the next day, transfected cells were lifted and seeded on a 96-well, white-opaque tissue culture-grade plate with a density of 0.04 x 10^4^ cells per well. The next day, action spectra for each MeOp mutant and WT were obtained using the TECAN Spark advanced multimode microplate reader. Before luminescence measurements, 10 μM 11CR and 5 μM aequorin were added to the well and incubated for 3 minutes before the baseline luminescence was taken. Next, luminescence data were collected in response to different wavelengths of light (ranging from 230-650 nm). Each well was only exposed to one light wavelength. Luminescence data were collected and normalized to the baseline to calculate the luminescent fold increase in response to each wavelength. Data were further processed, and action spectra were plotted using the Origin-pro data analysis software (OriginLab®).

### 3.6 Expression and purification of MeOp pigments and spectroscopy

Opsin expression and purification were performed as described previously^44^. Briefly, the full-length cDNAs expressing MeOp-WT and mutants were tagged with the monoclonal antibody rho 1D4 epitope sequence (ETSQVAPA) and were inserted into the pMT vector. Opsin expression vectors were transfected into HEK293S cells using the calcium-phosphate method. 11CR (final concentration of 2.7 μM) was added to the culture medium 24 hours after transfection and cultured in the dark for another 24 hours, according to the previous report^28^. Cells were harvested 48 hours post-transfection. The expressed proteins were incubated with an excess of 11CR overnight to reconstitute the pigment. Pigments were then extracted with 1% (w/v) dodecyl β-D-maltoside (DM) in HEPES buffer (pH 6.5) containing 140 mM NaCl and 3 mM MgCl_2_, bound to 1D4-agarose, washed with 0.02% DM in the HEPES buffer and eluted with the HEPES buffer containing 0.02% DM and 1D4 peptide. The absorption spectra of the opsin-based pigments were recorded at 4 °C using the V-750 UV-VIS Spectrophotometer (JASCO International). Blue light (440 nm) was supplied using A light source with a 440 nm interference filter to obtain the light spectrum.

### 3.7 Computational methods

ARM protocol was used for the fast and automatic generation of combined (QM/MM) models of rhodopsin-like receptors as previously described^41,39,54^ to predict opsin model absorption maxima as an average from the produced standard ten structural replicas. Briefly (see the cited references for details), the combination of the multi-configurational complete active space self-consistent field (CASSCF)^40,54^ method and the Amber force field is used to optimize the ground state geometries (single-root CASSCF(12,12)/6-31G*/Amber. The active space comprises the entire set of 12 electrons and 12 molecular orbitals describing the chromophore π-system). Excitation energies are then computed using multi-configurational second-order perturbation theory (CASPT2)^54^, using a three-roots stage average (SA) CASSCF(12,12)/6-31G* wavefunction as a reference.

The structure of the ARM-generated QM/MM model is also described briefly. The model is divided into three subsystems called Environment, Cavity, and Lys-QM. The protein environment (treated at MM level) has a backbone and side-chain atoms fixed at the X-ray (or homology model) structure. The effect of the protein environment is indirectly incorporated with the introduction of counterions (Cl− and/or Na+), which generate a model with neutral inner and outer surfaces. The relaxed chromophore cavity (treated at MM level) has fixed backbone atoms, but its side-chain atoms are free to relax. It contains all the amino acid residues around the retinal chromophore. ARM uses the CASTp server (http://sts.bioe.uic.edu/castp/index.html ?2cpk) to obtain the cavity residues list. Lys-QM subsystem comprises the atoms of the lysine (though C_δ_) to the entire retinal chromophore. All Lys-QM atoms are kept free to relax.

### 3.8 Statistical analysis

All experiments were repeated multiple times to test the reproducibility of the results. Statistical analysis and data plot generation were done using OriginPro software (OriginLab®). Results were analyzed from multiple cells on multiple days and represented as mean ± SD. The exact number of cells used in the analysis is given in respective figure legends. One-way ANOVA statistical tests were performed using OriginPro to determine the statistical significance between two or more populations of signaling responses. Tukey’s mean comparison test was performed at the p < 0.05 significance level for the one-way ANOVA statistical test.

## Acknowledgment

We acknowledge Dr. N. Gautam for providing us with plasmid DNA of various G protein subunits and for the RNA seq data. We thank the National Eye Institute for providing 11-*cis*-retinal. We thank Robert S. Molday (University of British Columbia) for supplying rho 1D4-producing hybridoma. We also thank Dinesh Kankanamge for their experimental assistance.

## Author contributions

D. W. And F.S. contributed equally to the study. D.W. conducted the action spectra, the majority of live cell imaging experiments, and the data analysis. F.S. performed all the computational analyses using modeller and ARM. L.P.G. developed the ARM protocol and F.F. generated the mMeOp homology model.M.K. T.S. and A.T. conducted the MeOp pigment purification and spectroscopic studies. S.P. conducted the Gγ9 translocation experiments and assisted in data analysis. K.G. performed the control experiments for the live cell imaging assay for bistability. W.T. made the mTq-PH construct. A.K., D.W., and M.O. conceptualized the project and wrote the manuscript.

## Data availability statement

The datasets used and/or analyzed during the current study are available from the corresponding authors upon reasonable request.

