## Supplementary Information for "Engineered blue-shifted melanopsins for subcellular optogenetics"

Running title: Spectral tuning of melanopsin

[1]Department of Chemistry, Saint Louis University, Saint Louis, MO 63103, USA
[2]Department of Biotechnology, Chemistry and Pharmacy, University of Siena, Siena, Italy
[3]Dulbecco Telethon Institute, Department of Life Sciences, University of Modena and Reggio Emilia, I-41125 Modena, Italy
[4]Department of Biology, Osaka Metropolitan University, O 3-3-138 Sugimoto, Sumiyoshi-Ku, Osaka, 558-8585, Japan
[5]The OMU Advanced Research Institute for Natural Science and Technology, Osaka Metropolitan University, Osaka, Japan
[6]Department of Biology, Siena Heights University, Adrian, MI, 49221, USA
[7]Department of Chemistry, Bowling Green State University, Bowling Green, OH, 43403, USA

[†]Authors contributed equally

**Keywords**: Melanopsin, Opn4, Optogenetics, Optical control, subcellular, Signaling, GPCR, G proteins

#### Figure S1

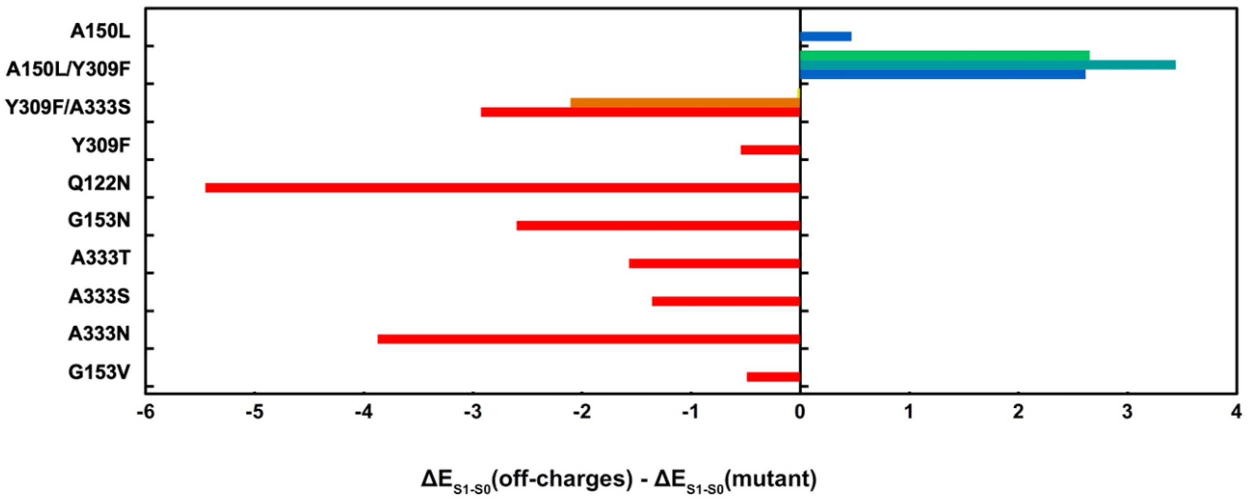


**Figure S1. Electrostatic contribution of the amino acid mutations in the active site of mMeOp. Red/blue bars indicate red/blue-shifts computed by zeroing the charges of each mutation.**

#### Figure S2

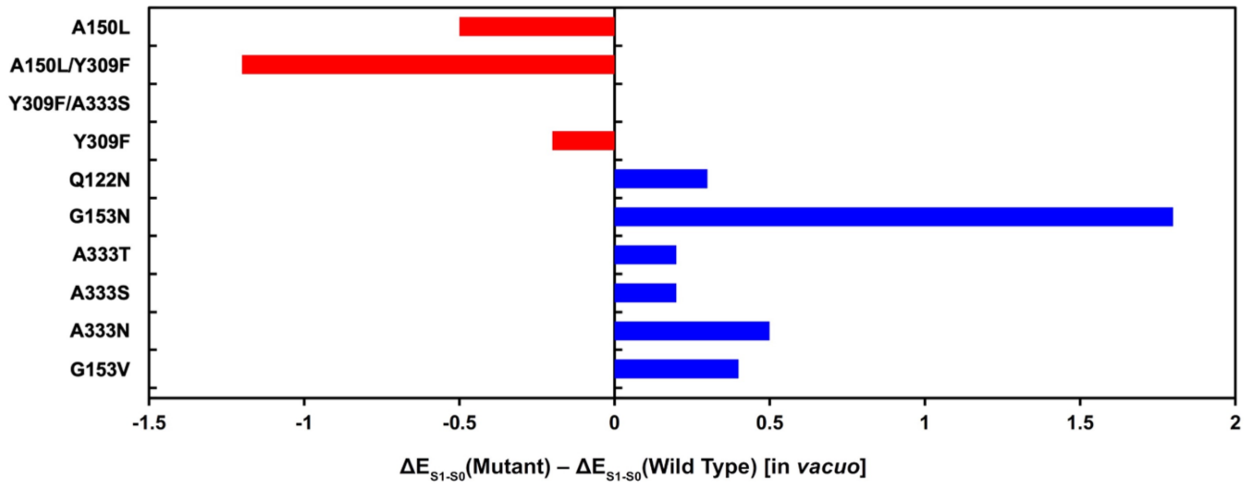


**Figure S2. Steric contribution of the amino acid mutations in the active site of mMeOp. Red/blue bars indicate red/blue-shifts as a consequence of steric interactions.**

### Figure S3

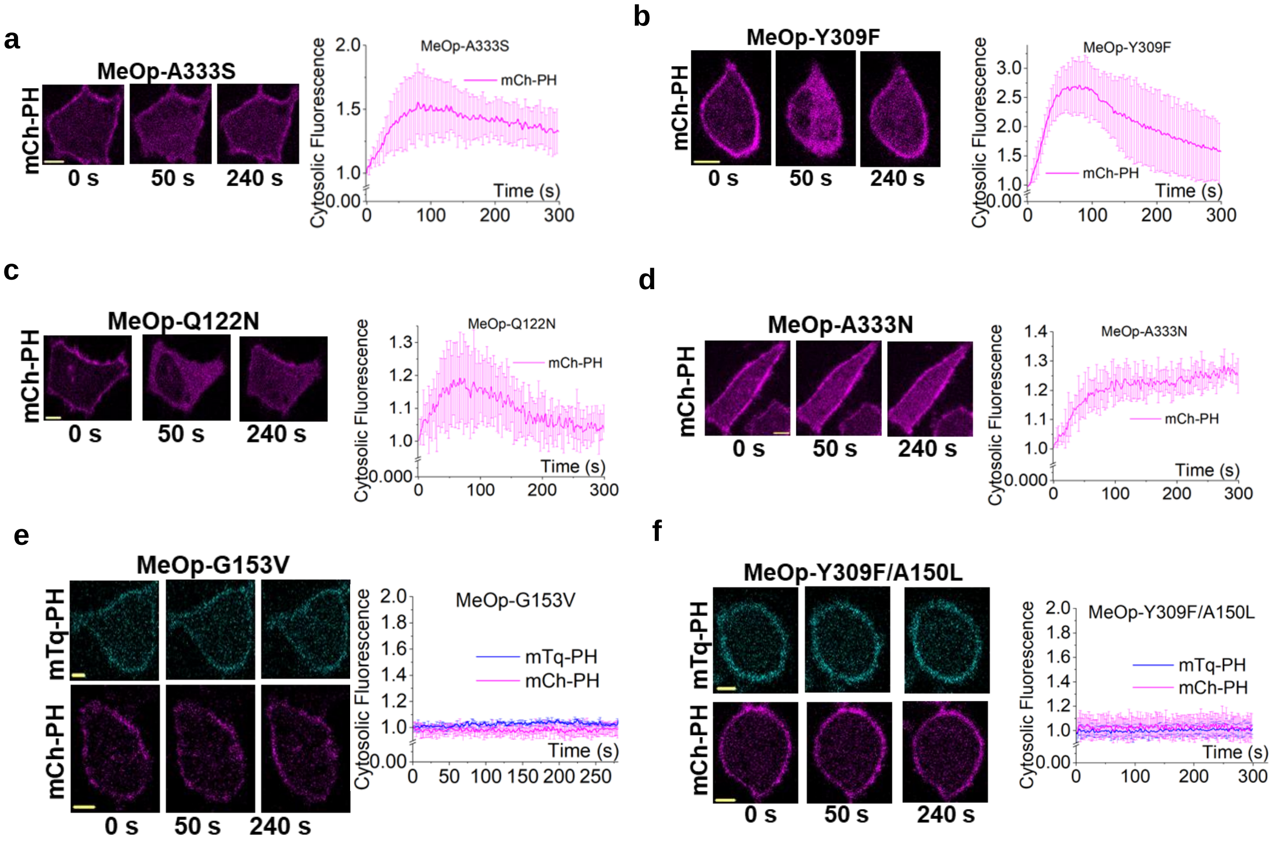


**Figure S3. a-d. MeOp mutants showed PIP2 hydrolysis with red light activation indicating inadequate blue shift. e-f. G153V and Y309F/A150L mutants showed no activity in red or blue light indicating a complete loss of activity due to the mutations.**

**Table S1:** Comparison between observed and computed vertical excitation energies, $\Delta E_{S1-S0}$ (kcal/mol) (maximum absorption wavelengths, $\lambda_{max}$ (nm)) for SqRh and mMeOp ARM models. The oscillator strength ($f_{osc}$), error between computational and experimental values (Error) and standard deviation (STD) are shown. The previously documented expected $\Delta E_{S1-S0}$ error produced by ARM models is a <3 kcal/mol blue-shifted error).

| Sample | Obs. $\Delta E_{S1-S0}$ ($\lambda_{max}$) | Comp. $\Delta E_{S1-S0}$ ($\lambda_{max}$) | $f_{osc}$ $S_0 \rightarrow S_1$ | Error | STD |
|---|---|---|---|---|---|
| SqRh | 58.5 (489) | 59.5 (480) | 0.82 | 1.0 | 0.20 |
| mMeOp | 61.2 (467) | 62.4 (457) | 0.77 | 1.2 | 0.48 |

**Table S2:** Comparison of the vertical excitation energies $\Delta E_{S1-S0}$ computed at the CASPT2/CASSCF(12,12)/6-31G*/Amber level of theory for the ARM models of SqRh and mMeOp. "Protein Environment" and "Vacuo" indicate a computation with the retinal chromophore inside and outside the opsin cavity respectively. In both cases the chromophore geometry is always that obtained inside the opsin cavity. All values are reported in kcal/mol. In all cases, the difference between SqRh and mMeOp excitation energies are reported inside the parenthesis.

| Sample | $\Delta E_{S1-S0}$ Protein Environment | $\Delta E_{S1-S0}$ *Vacuo* | Cavity Electrostatic Contribution |
|---|---|---|---|
| SqRh | 59.5 | 48.3 | 11.2 |
| mMeOp | 62.4 (+2.9) | 47.3 (-1.0) | +15.1 (+3.9) |

**Table S3: Comparison between observed and computed vertical excitation energies, ΔES1−S0 (kcal/mol) (maximum absorption wavelengths, λmax (nm)) for mutated mMeOp ARM models.** The oscillator strength (fosc), and error between computational and experimental values (Error) are shown.

| Sample | Obs. $\Delta E_{S1-S0}$ ($\lambda^a$max) | Comp. $\Delta E_{S1-S0}$ (λmax) | $f_{osc}$ $S_0 \rightarrow S_1$ | Error | Red light sensitivity |
|---|---|---|---|---|---|
| A333T | 67.9 (421) | 63.9 (447) | 0.76 | 4.0 | No |
| A150L | 69.6 (411) | 69.0 (415) | 0.52 | 0.6 | No |
| G153N | 68.2 (419) | 63.9 (447) | 0.77 | 4.3 | No |
| Y309F/ A333S | 68.0 (420) | 66.2 (433) | 0.69 | 1.8 | No |
| Y309F | No spectrum | 65.5 (436) | 0.71 | | Yes |
| Q122N | No spectrum | 64.2 (445) | 0.75 | | Yes |
| A333N | No spectrum | 63.2 (452) | 0.78 | | Yes |
| A333S | No spectrum | 63.4 (451) | 0.77 | | Yes |
| G153V | No spectrum | 62.9 (454) | 0.81 | | Lost all activity |
| A150L/ Y309F | No spectrum | 68.0 (420) | 0.64 | | Lost all activity |

**Table S4-a: One-way ANOVA statistics for PIP2 hydrolysis extent with MeOp WT and mutants, A333T, A150L, G153N, and Y309F/A333S.**

| **Descriptive Statistics- Gγ9 translocation attenuation extent** | | | | |
|---|---|---|---|---|
| | N Analysis | Mean | Standard Deviation | SE of Mean |
| WT | 10 | 1.08725 | 0.28135 | 0.08897 |
| A333T | 15 | 1.55376 | 0.40244 | 0.10391 |
| A150L | 19 | 1.06181 | 0.35102 | 0.08053 |
| G153N | 9 | 0.86266 | 0.46574 | 0.15525 |
| Y309F/A333S | 10 | 1.06115 | 0.51779 | 0.16374 |

**Table S4-b**

| **Overall ANOVA- Gγ9 translocation attenuation extent** | | | | | |
|---|---|---|---|---|---|
| | DF | Sum of Squares | Mean Square | F Value | Prob>F |
| Model | 4 | 3.45329 | 0.86332 | 5.35772 | 9.66944E-4 |
| Error | 58 | 9.34589 | 0.16114 | | |
| Total | 62 | 12.79918 | | | |

At the 0.05 level, the population means are **significantly** different.

**Table S5-a: One-way ANOVA statistics for Gγ9 translocation extent with MeOp mutants, A333T, A150L, G153N, and Y309F/A333S.**

| **Descriptive Statistics- Gγ9 translocation attenuation extent** | | | | |
|---|---|---|---|---|
| | N Analysis | Mean | Standard Deviation | SE of Mean |
| WT | 11 | 0.22979 | 0.08615 | 0.02597 |
| A333T | 13 | 0.07707 | 0.02 | 0.00555 |
| A150L | 12 | 0.14753 | 0.05039 | 0.01455 |
| G153N | 11 | 0.03943 | 0.0144 | 0.00434 |
| Y309F/A333S | 9 | 0.11503 | 0.05712 | 0.01904 |

**Table S5-b**

| **Overall ANOVA- Gγ9 translocation attenuation extent** | | | | | |
|---|---|---|---|---|---|
| | DF | Sum of Squares | Mean Square | F Value | Prob>F |
| Model | 4 | 0.23723 | 0.05931 | 22.38526 | 1.02508E-10 |
| Error | 51 | 0.13512 | 0.00265 | | |
| Total | 55 | 0.37236 | | | |

At the 0.05 level, the population means are **significantly** different.

**Table S6-a: One-way ANOVA statistics for Gγ9 translocation attenuation with blue opsin and lamprey parapinopsin activation with 50 nM 11-cis-retinal**

| **Descriptive Statistics- Gγ9 translocation attenuation extent** | | | | |
|---|---|---|---|---|
| | N Analysis | Mean | Standard Deviation | SE of Mean |
| Lamprey parapinopsin | 12 | 0 | -5.77611 | 7.72238 |
| Blue opsin | 13 | 0 | 67.78169 | 10.65141 |

**Table S6-b**

| **Overall ANOVA- Gγ9 translocation attenuation extent** | | | | | |
|---|---|---|---|---|---|
| | DF | Sum of Squares | Mean Square | F Value | Prob>F |
| Model | 1 | 33763.08228 | 33763.08228 | 384.92321 | 7.34759E-16 |
| Error | 23 | 2017.41771 | 87.71381 | | |
| Total | 24 | 35780.49999 | | | |

At the 0.05 level, the population means are **significantly** different.

**Table S7-a: One-way ANOVA statistics for blue opsin-induced Gγ9 translocation attenuation with 50 nM and 10 μM 11CR**

| **Descriptive Statistics- Gγ9 translocation attenuation extent** | | | | |
|---|---|---|---|---|
| | N Analysis | Mean | Standard Deviation | SE of Mean |
| 50 nM 11CR | 8 | 52.88668 | 10.36146 | 3.66333 |
| 10 μM 11CR | 8 | 1.2713 | 3.80472 | 1.34517 |

**Table S7-b**

| **Overall ANOVA- Gγ9 translocation attenuation extent** | | | | | |
|---|---|---|---|---|---|
| | DF | Sum of Squares | Mean Square | F Value | Prob>F |
| Model | 1 | 10656.58876 | 10656.58876 | 174.93367 | 2.65722E-9 |
| Error | 14 | 852.85034 | 60.91788 | | |
| Total | 15 | 11509.4391 | | | |

At the 0.05 level, the population means are **significantly** different.
